# cfMethDB: a comprehensive cfDNA methylation data resource for cancer biomarkers

**DOI:** 10.1101/2025.03.19.644046

**Authors:** Yuanhui Sun, Zhixian Zhu, Qiangwei Zhou, Zhe Wang, Yuying Hou, Xionghui Zhou, Guoliang Li

**Author notes:** Equal contribution (Yuanhui Sun, Zhixian Zhu, Qiangwei Zhou). Corresponding author. (Li G).

## Abstract

Cancer is a major global health threat, and early detection of cancer is crucial for improving patient outcomes. DNA methylation in circulating cell-free DNA (cfDNA) has emerged as a promising biomarker for non-invasive cancer diagnosis. However, the integration and utilization of existing cfDNA methylation data have been limited, hindering comprehensive research efforts, especially in the discovery of cfDNA methylation biomarkers. To address this challenge, we introduce cfMethDB, a comprehensive database dedicated to cfDNA methylation in cancer that encompasses 4828 publicly available datasets. Through standardized analysis, we identified 1,048,770 differentially methylated cytosines (DMCs) as candidate biomarkers across seven cancer types. With cfMethDB, we not only identified known cfDNA methylation biomarkers, but also discovered several genes, such as *ZIC4*, that could be novel biomarkers. Moreover, cfMethDB offers a suite of user-friendly tools, including biomarker evaluation, pan-cancer search and end motif analysis. We hope that cfMethDB will serve as a valuable platform for the discovery of novel cancer cfDNA methylation biomarkers and will facilitate cancer research and clinical applications. cfMethDB is publicly available at: https://cfmethdb.hzau.edu.cn/home.

## Introduction

Cancer is a major threat to human health, with nearly 20 million new cases and 9.7 million deaths reported in 2022 [1]. Early detection of cancer is a viable strategy for improving outcomes in terms of population health [2]. For instance, delays in breast cancer treatment, particularly those longer than three months, are associated with advanced-stage diagnosis and reduced survival rates [3]. In the THUNDER study, the interception model projected that, compared to usual care, 38.7% to 46.4% of patients who had been diagnosed at late stages could be identified at earlier stages using the early diagnosis method [4].

For early cancer diagnosis, minimally invasive sampling methods, such as collecting cell-free DNA (cfDNA) from the plasma, are preferred [5]. cfDNA is released by dead cells into the bloodstream [6], offering a new method for non-invasive cancer diagnosis [7]. Many cancer-associated molecular features have been detected in the cfDNA of patients with various cancer types [8]. Among those features, cfDNA methylation has emerged as the most promising signal for cancer detection [2]. Recently, many studies utilize cfDNA methylation signals for early diagnosis of various cancers. For example, methylation of *SEPT9* can be used to detect majority of colorectal cancer patients at all stages [9]. Similarly, the combination of *HOXA9* and *HIC1* methylation serves as an effective diagnostic biomarker for early ovarian cancer screening [10]. Promoter methylation of *GSTP1* is highly specific for diagnosing prostate cancer [11]. Many methylation biomarkers have also been identified in other cancer types, including lung cancer [12], liver cancer [13] and breast cancer [14].

With the blooming development of cfDNA methylation diagnosis, a vast amount of cfDNA methylation data has been generated. Although some databases contain cfDNA methylation data, effectively integrating and utilizing these data remains challenging. The CFEA database encompasses three widely recognized epigenetic modifications information on cfDNA; however, the cfDNA methylation data is limited and has not been updated recently [15]. cfOmics provides multi-omics liquid biopsy data, including cfDNA and cfRNA; however, this database does not focus primarily on cfDNA methylation data, and lacks DNA methylation information at single-base resolution [16]. These limitations hinder the comprehensive exploration of cfDNA methylation across the whole genome and particularly effect the discovery of cfDNA methylation biomarker for cancer diagnosis. In addition, the characteristics of cfDNA fragments, such as fragment sizes and end motifs, are crucial for cfDNA analysis [17, 18]. Incorporating these fragment features into the cfDNA database is essential for obtaining a deeper understanding of cfDNA.

In this work, we introduce cfMethDB (https://cfmethdb.hzau.edu.cn/home), a comprehensive cfDNA methylation data resource for cancer biomarkers, which includes 4828 publicly available cfDNA methylation datasets. Using a standardized analysis pipeline, we identified 1,048,770 differentially methylated cytosines (DMCs) from cfDNA methylation data as candidate biomarkers across 7 types of cancer. With cfMethDB, users can explore these biomarkers across the entire genomic landscape and are not limited to specific regions, such as genes or promoters. cfMethDB provides detailed information on DMCs and a wide range of functions, including region searching, pan-cancer DMC searching and diagnostic evaluation tool, to help researchers explore candidate methylation biomarkers. Moreover, cfMethDB offers information on the fragmentation features of cfDNA, assisting researchers in exploring the intrinsic properties of cfDNA.

cfMethDB is the first database to provide cancer cfDNA methylation biomarkers with precise genomic location information. With cfMethDB, researchers can easily explore the differences in cfDNA methylation in cancer, facilitating clinical diagnosis and treatment.

## Data collection and processing

### Data collection and processing

We collected cfDNA methylation data from the Sequence Read Archive (SRA) database [19] and Genome Sequence Archive (GSA) database [20] up to June 2024. A total of 7530 datasets were downloaded initially (**Figure 1A**).

**Figure 1.**
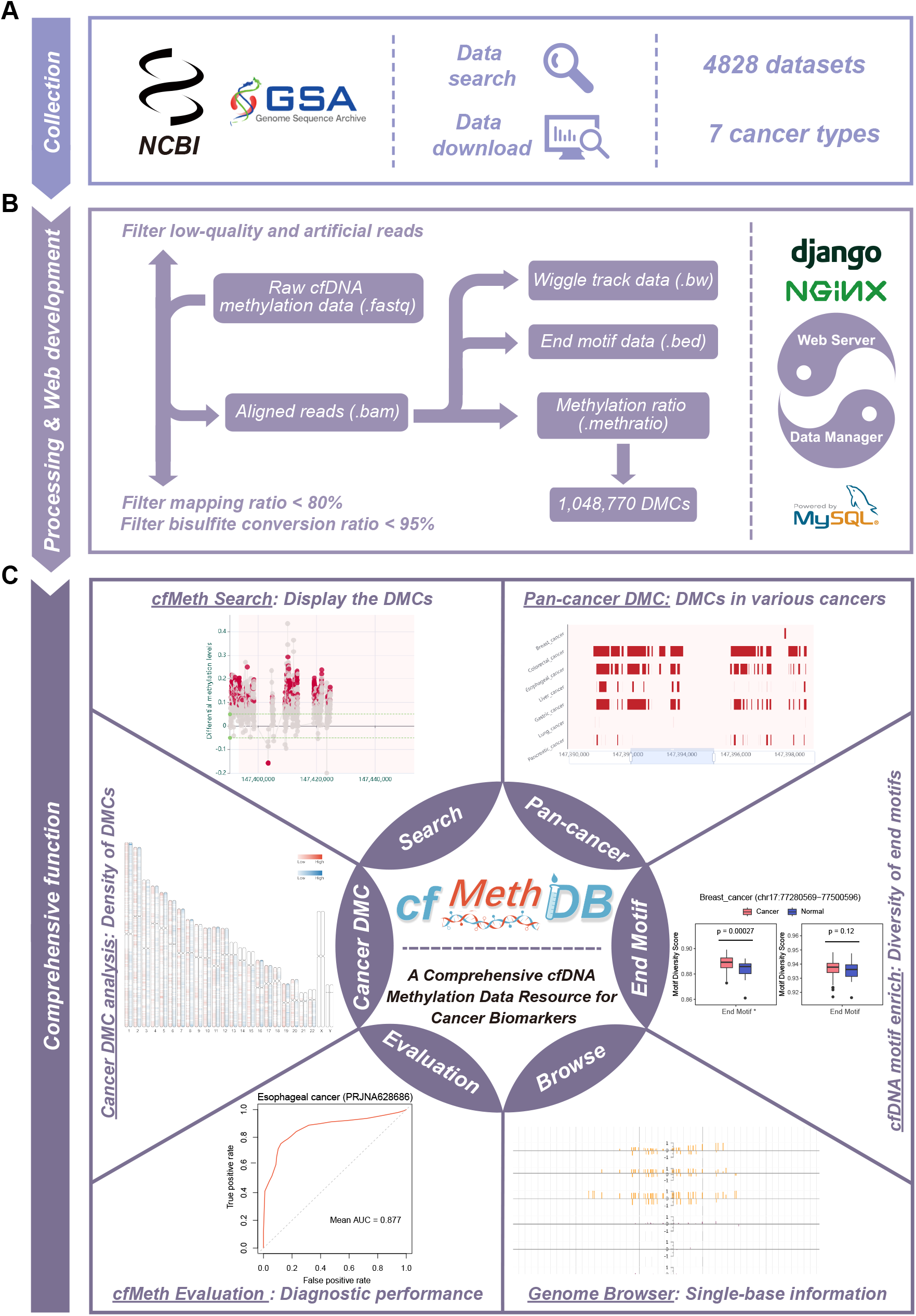
Workflow of the cfMethDB database. **A**. Collection of cfDNA methylation datasets from SRA and GSA databases. **B**. Data processing and web development for cfMethDB. **C**. Comprehensive functional and analysis modules in cfMethDB.

The collected DNA methylation sequencing data were generated from various methods, including WGBS, RRBS, and padlock-based methods. The only different in data analysis among these sequencing methods lies in the preprocessing stage. Fastp [21] was used to trim the low-quality and artificial reads with default parameters. Specifically, we used Trim Galore with the ‘*-rrbs*’ parameter, which identifies sequences that were subjected to adapter contamination and removes another 2 bp from the end of reads to account for technical bias introduced by the RRBS technique [22]. BatMeth2 [23] was subsequently used to map the trimmed reads to the human reference genome (hg38) and the mapping ratio was calculated using SAMTools [24]. Datasets with mapping ratio below 80% were filtered out. After mapping, the reads with the alignment quality score below 30 and the coverage of each cytosines less than 2 were discarded. The ‘Calmeth’ module in BatMeth2 was used to calculate the DNA methylation level of each cytosine (**Figure 1B**). The bisulfite conversion rate was evaluated by calculating the CHG methylation level as described previously [25] and datasets with the conversion rate lower than 95% were excluded from downstream analysis. Finally, 4828 datasets were retained for further analysis and the number of CpG sites per samples was displayed in **Figure S1A**.

### Collection of biomarker genes from literature

Initially, we retrieved abstracts of published literature from PubMed and extracted sentences related to DNA methylation, various cancer types, and genes. After manual curation, we compiled a total of 2611 biomarker records, including 87 DNA methylation biomarker genes in cfDNA and 697 genes in tissue, respectively. In total, cfMethDB includes 729 unique genes in the curated biomarker records. We utilized word cloud plots and tables for intuitive visualization of these biomarker records.

### Differentially methylated cytosine (DMC) analysis

To identify the DMCs across the whole genome for each cancer type, we used SMART2 software [26] in DMC mode with default parameters (Figure 1B). The DMCs with an absolute methylation difference greater than 0.05 and a p-value less than 0.01 were retained for downstream analysis. The ‘ChIPseeker’ R package [27] was used to annotate these DMCs. The DMCs included in cfMethDB covered most of the known DNA methylation biomarkers, including approximately 94% (686 over 729) of the DNA methylation biomarkers previously reported in the literature, as well as all 37 DNA methylation biomarkers approved by the National Medical Products Administration (NMPA) (**Table S1**).

### cfDNA fragmentomics analysis

cfDNA fragmentation patterns are also promising biomarkers for cancer detection. For example, the frequency of end motifs can effectively differentiate cancer samples from normal samples [17]. In this work, the cfDNA fragment sizes were calculated with the ‘CollectInsertSizeMetrics’ module in GATK [28]. The distribution of cfDNA fragment lengths across different projects was illustrated in **Figure S1B**. For end motifs, we used BEDTools [29] to extract the first 4 bases from the 5’ ends of the fragments. We also introduced ‘U’ to indicate unmethylated cytosines to distinguish the methylation status of the cytosines. The end motif frequency was calculated by dividing the total number of reads in the region by the number of occurrences count of each motif. The motif diversity score was computed using the methodology outlined in a previous study [17].

### Diagnostic analysis of methylation biomarkers

In cfMethDB, users can select the cancer type and project of interest, and the DNA methylation levels are extracts from the queried regions. Any queried regions with missing values more than 20% were excluded from further analysis. For the remaining queried regions, missing values were imputed using the ‘na_interpolation’ function of the ‘imputeTS’ R package [30], which employs a linear model. Then, a logistic regression-based diagnostic model was constructed with the ‘caret’ R package [31] on the whole datasets of selected project, with a twenty times five-fold cross-validation process.

### Database implementation

cfMethDB includes several search and analysis functions complemented by a genome browser powered by JBrowse [32]. The data within the cfMethDB database were organized by MySQL, and the web interface was constructed by Nginx and the Django framework, delivering a user-friendly experience for data exploration (**Figure 1B, C**).

## Database content and usage

### Overview of cfMethDB

cfMethDB is an integrative resource for cfDNA methylation data (**Figure 2A**). The current version of cfMethDB includes 4828 datasets from 7 different cancer types, with a total of 1,048,770 DMCs (**Figure 2B, C; Table S2, S3**). With a focus on the identified genome-wide DMCs, cfMethDB serves as a user-friendly platform providing a variety of function modules (Figure 1C):

**Figure 2.**
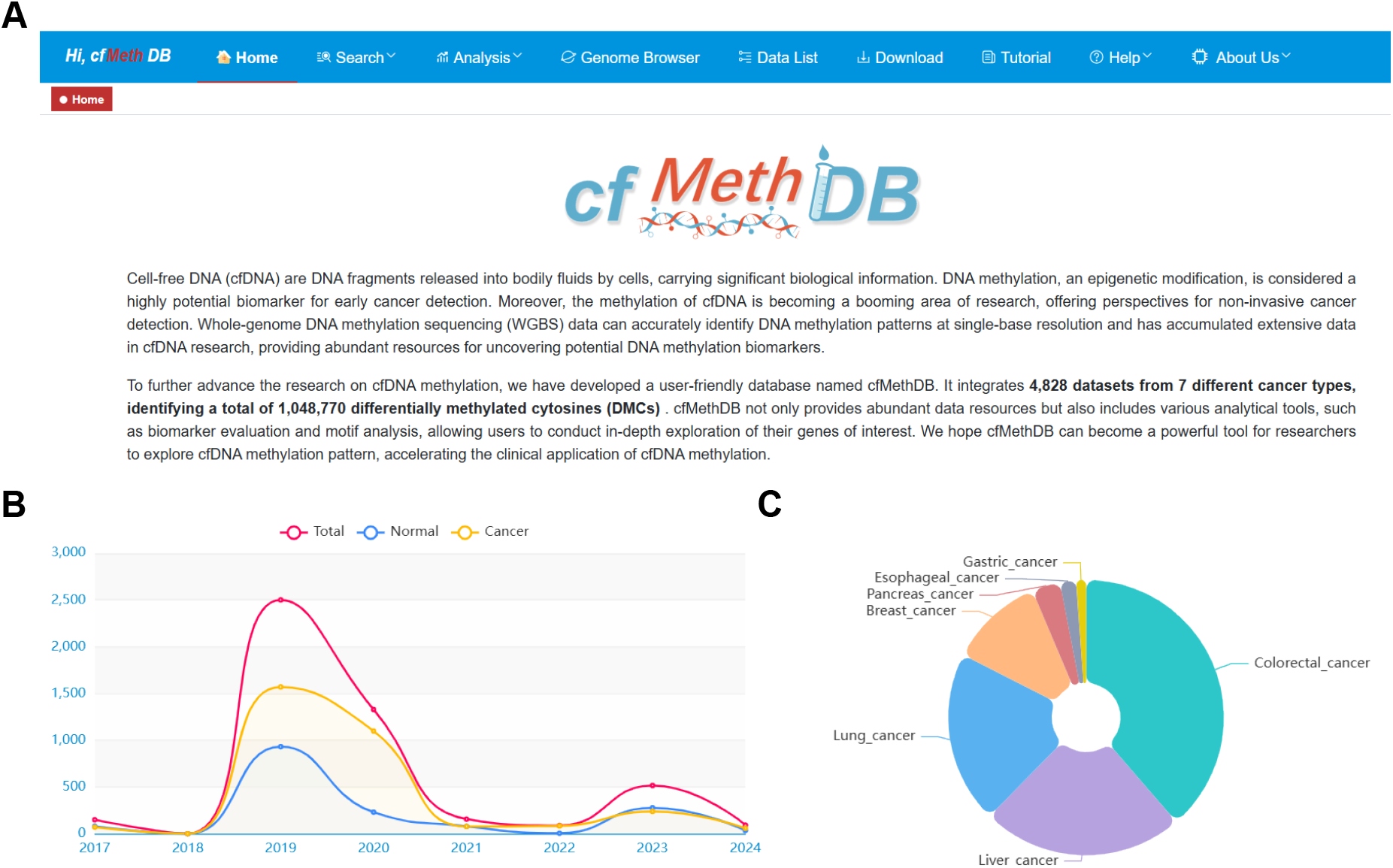
Overview of the cfMethDB database. **A**. Homepage and brief introduction about cfMethDB. **B**. Statistics of cfDNA methylation datasets in cfMethDB per year. **C**. Proportion of sample groups corresponding to cfDNA methylation datasets collected in cfMethDB.

(i) **Search** – This retrieval module allowing users to explore the detailed DMC information on queried genes (*cfMeth Gene Search*) or regions (*cfMeth Region Search*) in each cancer type. In addition, this module enables users to search for DNA methylation biomarker genes reported in the literature (*Literature Search*). cfMethDB also offers APIs that enable users to conveniently query *cfMeth Search* results, which can be incorporated into other databases or software packages.
(ii) **Analysis** – This module includes four online tools. The module provides detailed information on DMCs within a specific type of cancer (*Cancer DMC analysis*), as well as across various cancer types (*Pan-cancer DMC*). The module can also evaluate the diagnostic value of a series of user-specified genomic regions in a particular cancer type (*Marker evaluation*) and analyze the 5’ end motif patterns of genomic region (*End motif analysis*). The data used for the Analysis module are presented in **Table S4**.
(iii) **Genome Browser** – This module enables users to explore DNA methylation status at a single-base resolution.
(iv) **DataList** and **Download** – These modules provide access to cfDNA methylation dataset information and DMC result files for browsing and downloading.
(v) **Tutorial, Help** and **About Us** – These modules provide users with detailed documentation and tutorial.

### Function modules in cfMethDB

#### Gene search

The ‘cfMeth Gene Search’ module allows users to retrieve detailed DMC information for a particular cancer type. The search results include: (i) basic information about the queried gene; (ii) a distribution plot of DMCs annotated to the queried gene; (iii) detailed information on DMRs derived from the MethMarkerDB database [33] for the queried gene; (iv) gene expression information of the queried gene across different tissues from the GEPIA2 database [34].

Here, we use the Syndecan 2 (*SDC2*) gene as an example to demonstrate the ‘cfMeth Gene Search’ function and to showcase the credibility of the results within cfMethDB. The *SDC2* gene encodes a member of the heparan sulfate proteoglycans family, and plays diverse roles in cell adhesion and cell communication [35, 36]. Previous studies have demonstrated that hypermethylation of the *SDC2* promoter is a common epigenetic alteration in the development of colorectal cancer, as observed in both tissue and serum samples [37, 38]. Moreover, the *SDC2* gene is used in an early detection kit for colorectal cancer that was approved by the NMPA (**Table S1**).

On the search page, users can specify the cancer type and the gene of interest, such as the *SDC2* gene in colorectal cancer (**Figure 3A**). The distribution plot of DMCs for the *SDC2* gene shows a hypermethylation region in the promoter region. Detailed information on these DMCs is displayed in a table (**Figure 3B, C**). In addition, the genome browser can display single-base DNA methylation information in the hypermethylation region of the *SDC2* gene (**Figure 3D**). Furthermore, gene expression data reveals the lower gene expression levels in colorectal cancer samples than in normal samples (**Figure 3E**), suggesting a potential correlation between hypermethylation and gene silencing.

**Figure 3.**
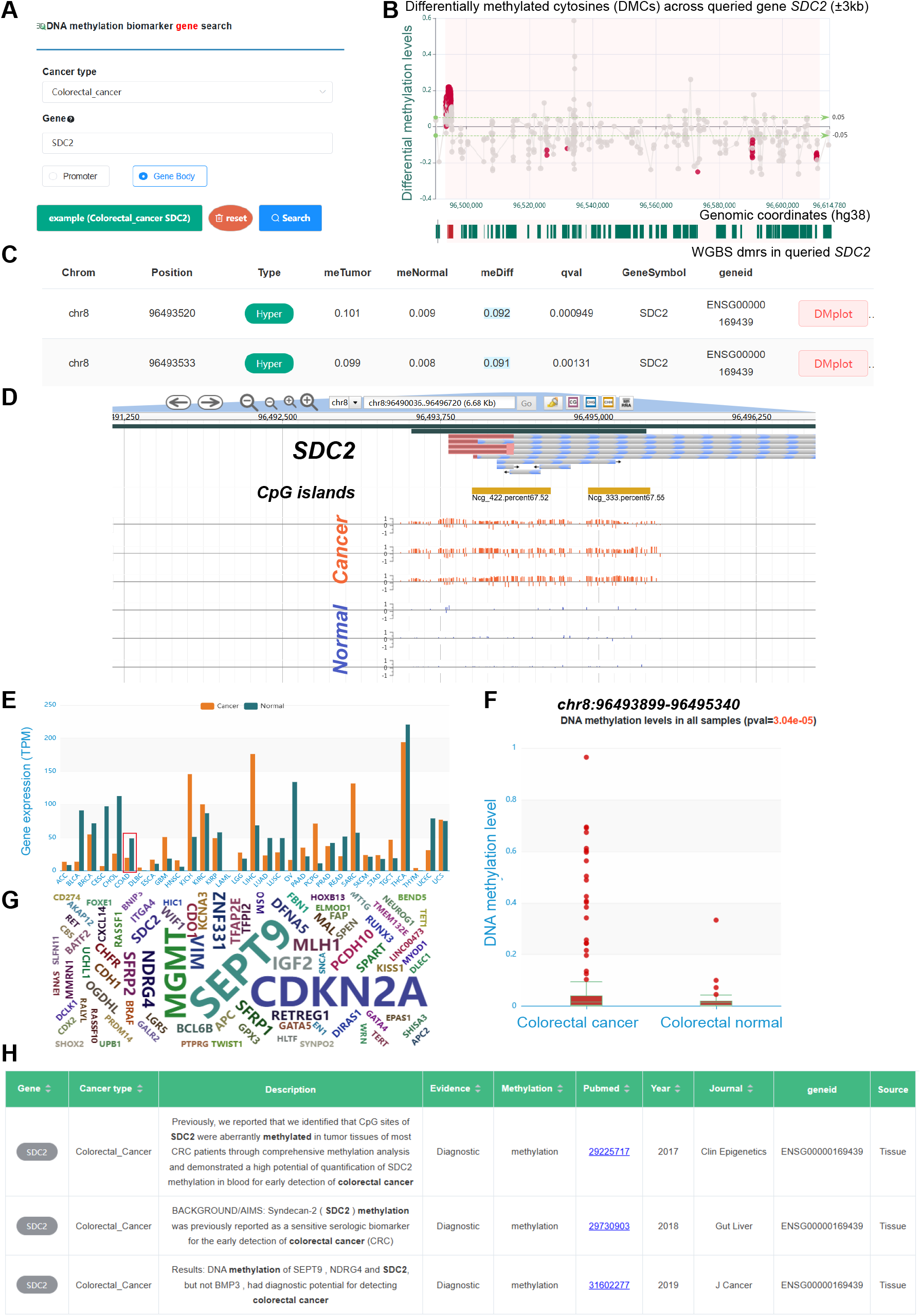
Search Modules in cfMethDB database. **A**. A screenshot of the gene search page. **B**. Distribution of DMCs across a 3 kb region upstream and downstream of the *SDC2* gene in colorectal cancer. **C**. Detailed information about the DMCs displayed in Figure 3B. **D**. A screenshot of genome browser showing the DNA methylation levels around the promoter region of *SDC2* in colorectal cancer. **E**. Expression levels of *SDC2* from the GEPIA2 database. **F**. DNA methylation levels of *SDC2* promoter regions (chr8:96,493,899-96,495,340) in colorectal cancer and normal samples. **G**. Word cloud showing the previously reported biomarker genes associated with colorectal cancer. **H**. Detailed records of the reported DNA methylation biomarkers of the *SDC2* gene.

#### Region search

Similar to ‘cfMeth Gene Search’ module, the ‘cfMeth Region Search’ module allows users to search for DMCs within genomic regions of interest for a specific cancer type. The search results include the detailed information about genes and DMCs in the queried region. Users can also access DNA methylation levels for the queried genomic region in each cancer type. For example, users can access the methylation level of the region (chr8:96,493,899-96,495,340) in colorectal cancer and normal samples (**Figure 3F**), enabling direct comparisons of the methylation levels between cancer and normal groups.

#### Literature search

In the ‘Literature Search’ module, users can select a specific cancer type, such as colorectal cancer. A word cloud plot is generated to display the reported DNA methylation biomarkers for that cancer type; for example, the *SEPT9, CDKN2A*, and *SDC2* genes in colorectal cancer (**Figure 3G**). Moreover, users can search for a specific gene of interest, such as *SDC2*, to obtain detailed records of this biomarker in tabular format (**Figure 3H**).

#### Genome browser

JBrowse is a convenient and user-friendly platform for browsing single-base DNA methylation levels. Users can select different samples to explore the DNA methylation level of a specific gene or region. The genome browser indicates that there are two CpG islands in the *SDC2* promoter region that exhibit a hypermethylation pattern in colorectal cancer compared with normal samples (Figure 3D). Moreover, JBrowse includes several plugins, such as ‘Share’ and ‘ScreenShot’ [32]. The ‘Share’ plugin allows users to generate a shareable URL for track information, and the ‘ScreenShot’ plugin enables users to save the screenshots in various format, such as PNG, JPG and PDF. Furthermore, JBrowse supports the upload of their local data for browsing, including genomic repeat regions, regulatory elements and sequencing signals in bigWig or BED formats.

#### Cancer DMC analysis

To better understand the distribution patterns of DMCs across the genome and to compare these patterns among different cancers, we integrated the DMCs detected in various cancer types and annotated them with genomic features. In the ‘Cancer DMC analysis’ module, users can browse and query the results of all DMCs for various types of cancer. **Figure 4A** displayed the distribution of hypermethylated and hypomethylated DMCs across each chromosome in esophageal cancer. The genomic annotation of these DMCs reveals distinct distribution patterns: hypermethylated DMCs are predominantly enriched in promoter regions, whereas hypomethylated DMCs are enrich in intronic and intergenic regions (**Figure 4B**). Interestingly, we identified four genomic regions characterized by a high density of hypermethylated DMCs across all cancer types (**Figure S2**). Notably, *HOXD* genes (chr2:176 Mb–176.5 Mb), *HOXA* genes (chr7:27 Mb–27.5 Mb), *PCDH* genes (chr5:141 Mb–141.5 Mb) and *ZIC1/ZIC4* genes (chr3:147 Mb–147.5 Mb) are located in these regions. Previous studies have reported that epigenetic dysregulation of clustered *PCDH* and *HOX* genes can serve as powerful diagnostic biomarkers [39, 40]. Overall, these results suggest that cfMethDB can serve as a powerful tool for potential biomarker discovery.

**Figure 4.**
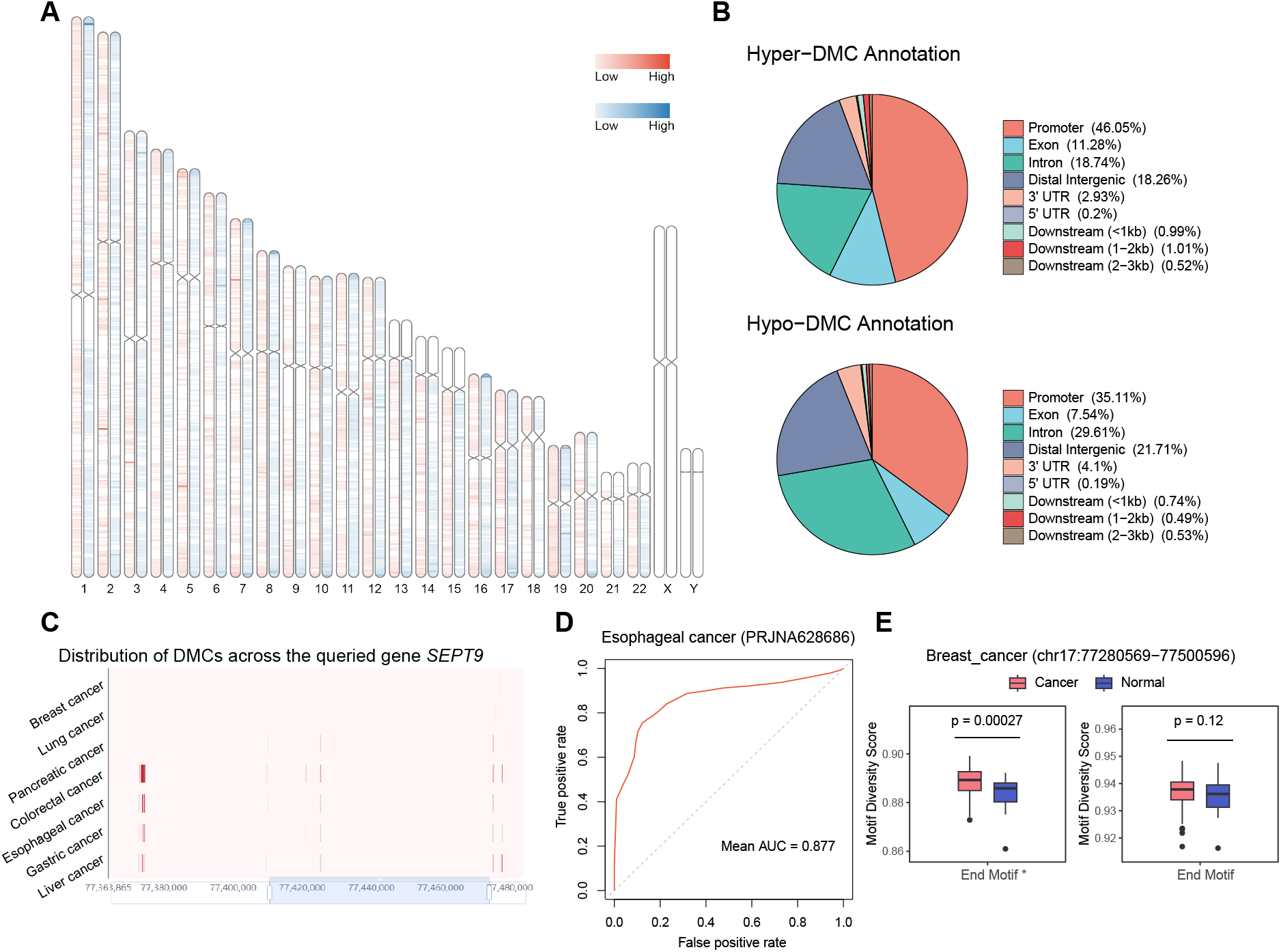
Function Modules in cfMethDB database. **A**. Chromosome ideogram plots displaying the density of DMCs in esophageal cancer. **B**. Genomic annotations for hypermethylated DMCs and hypomethylated DMCs in esophageal cancer, respectively. **C**. Distribution of DMCs in the *SEPT9* gene in various cancers. **D**. ROC curve for the *SEPT9* gene (using chr17:77,371,716 – 77,374,859 and chr17: chr17:77,428,290–77,431,433 as input regions) in classifying esophageal cancer and normal samples. **E**. Motif diversity score of breast cancer and normal samples. End Motif *: 625 end motifs with methylation status (‘U’ represents unmethylated cytosines). End Motif: 256 end motifs without methylation status. P values are calculated by Wilcoxon rank sum test.

#### Pan-cancer DMC analysis

Some studies have investigated pan-cancer DNA methylation patterns in tissue samples [33, 41, 42], but whether similar patterns are present in cfDNA remains unclear. To investigate pan-cancer patterns in cfDNA, we introduced the ‘Pan-cancer DMC’ module, which enables users to uncover potential pan-cancer methylation biomarkers. The *SIX6* and *SOX11* genes, which have been reported as promising pan-cancer biomarkers according to tissue data [33, 43], have also been found to be promising pan-cancer biomarkers in cfDNA (**Figure S3**). Additionally, through cfMethDB, we revealed that hypermethylation of the *SEPT9* gene could serve as a potential pan-cancer biomarker (**Figure 4C**). In essence, the ‘Pan-cancer DMC’ module facilitates the exploration of pan-cancer methylation patterns in cfDNA.

#### Biomarker evaluation

To assess the clinical application value of potential genomic regions effectively, we developed the ‘Marker evaluation’ module. This module enables users to evaluate the diagnostic performance of genomic regions of interest. Recently, the *SEPT9* gene was approved by the NMPA as a biomarker for esophageal cancer detection (**Table S1**). As expected, our results demonstrated that the diagnostic performance of the *SEPT9* gene in esophageal cancer is highly promising (**Figure 4D**). Furthermore, the ‘Marker evaluation’ module allows users to retrieve the DNA sequences of queried genomic regions, facilitating the design of primers using online tools.

#### End motif analysis

Recent studies have demonstrated that end motifs enable to reveal hallmarks of the non-random fragmentation process of cfDNA. For example, deoxyribonuclease 1L3 (*DNASE1L3*) might be linked to the generation of CCCA end motif, whereas deoxyribonuclease 1 (*DNASE1*) might be associated with the generation of the TGTG end motif [44, 45]. The diversity of end motifs suggests that these motifs can be new biomarkers for cancer detection in the emerging field of fragmentomics [17]. Despite the importance of end motifs, there are a limited number of web servers capable of analyzing the end motif of cfDNA. Jiang et al. demonstrated that the differential end motifs identified in WGS were also detectable in WGBS and there was a high correlation between the motif diversity scores from WGBS and WGS [17]. Given that fragmentation patterns may vary across different regions of the genome [46, 47], we developed the ‘End motif analysis’ module to enable users to explore the end motif pattern of genes or genomic regions of interest. For example, the motif diversity score of the *SEPT9* gene was significantly greater in breast cancer samples than in normal samples (**Figure 4E**), indicating that there was a wide variety of plasma DNA molecules with different end motifs in plasma from breast cancer samples. Interestingly, we also noticed that the difference in the motif diversity score between cancer and normal samples increased when the methylation status of the cytosine within the end motif was considered. A similar pattern was observed across the whole genome in different projects (**Figure S4**).

### Application of cfMethDB

Here, an example of how to use cfMethDB to discover potential biomarkers is presented. As we have described, users can identify potential biomarkers. The ‘Cancer DMC analysis’ module has shown that there is a high density of hypermethylated DMCs around the *ZIC1* and *ZIC4* genes. Using the ‘Pan-cancer DMC’ module, we identified several hypermethylated regions of the *ZIC4* gene that were consistent across various cancer types in cfDNA (**Figure 5A**). To further explore this gene, we utilized the ‘cfMeth Gene Search’ module to search for DMCs in liver cancer. The DMC distribution plot reveals that a high density of DMCs in the *ZIC4* gene, and the distribution of DMCs in the *ZIC4* gene is similar to the distribution of DMRs derived from tissue data (**Figure 5B**). Next, we used the ‘Genome browser’ module to display more detailed DNA methylation information for individual samples. It is evident that the *ZIC4* gene contains a hypermethylated region, particularly within a promoter for shorter transcripts. (**Figure 5C**). Furthermore, we found that the DNA methylation level of the *ZIC4* gene can effectively distinguish liver cancer plasma samples from normal plasma samples (**Figure 5D**). Overall, our analysis indicates that *ZIC4* could be a potential biomarker for liver cancer diagnosis.

**Figure 5.**
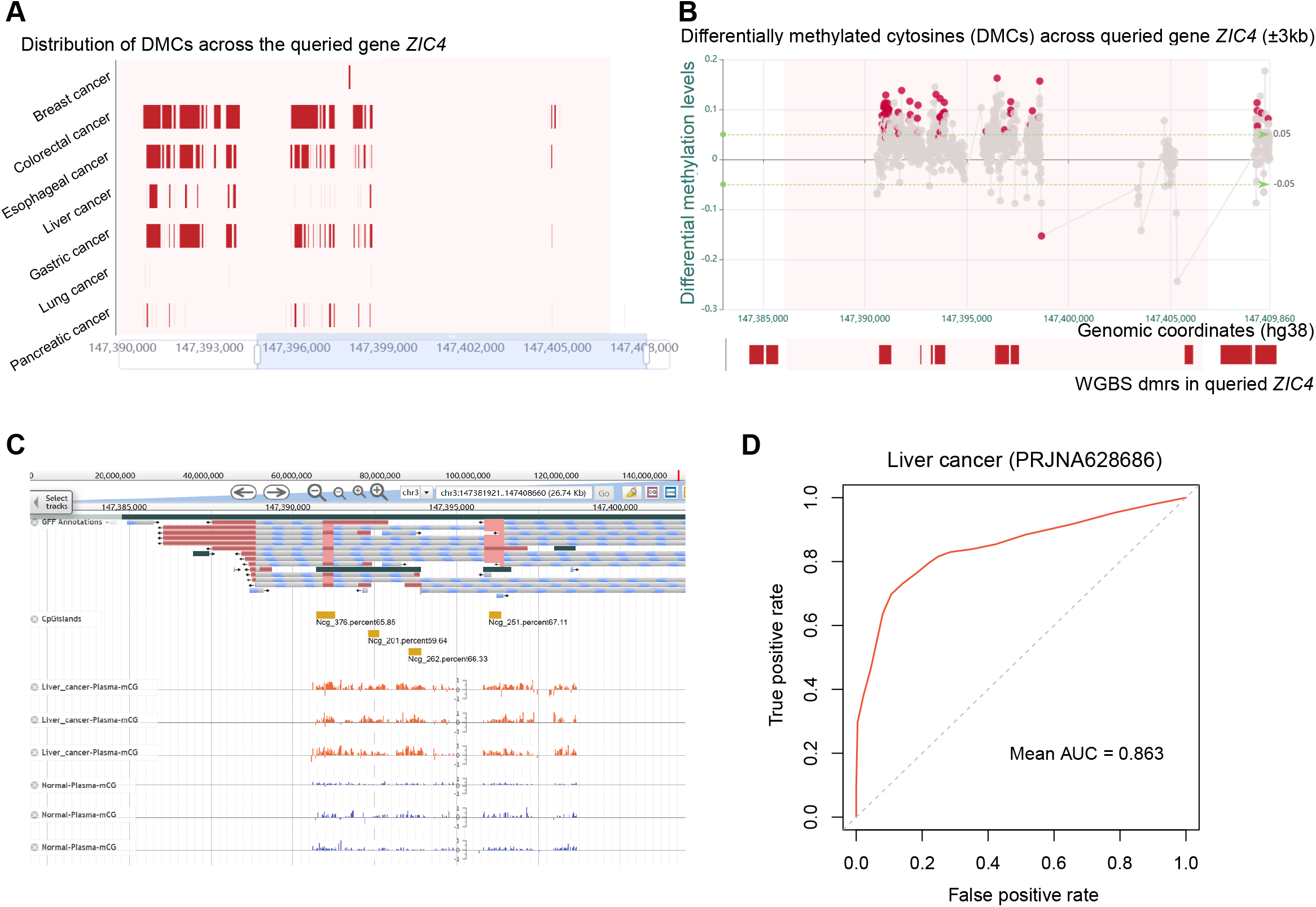
A case study in cfMethDB database. **A**. Distribution of DMCs in the *ZIC4* gene in various cancers. **B**. Distribution of DMCs across a 3 kb region upstream and downstream of the *ZIC4* gene in liver cancer. **C**. A screenshot of genome browser showing the DNA methylation levels around the promoter region of *ZIC4* in liver cancer. **D**. ROC curve for *ZIC4* gene (using chr3:147,391,246 – 147,391,766; chr3:147,391,970 – 147,392,702 and chr3:147,402,686–147,403,518 as input regions) in classifying liver cancer plasma samples and normal samples.

### Download and other features in cfMethDB

The ‘Download’ module provides carefully curated biomarker information from published studies and identified DMCs for various cancer types. Additionally, cfMethDB allows users to download cfDNA methylation via the ‘DataList’ module, with which users can search for and access samples of interest. For each sample, cfMethDB provides detailed information, including the SRA ID, sample source, PubMed ID and bisulfite conversion rate (**Figure S5A**). The methylation levels and data quality of each sample, including the coverage and the depth of sequencing data and the distribution of cfDNA fragments, are also displayed (**Figure S5B-E**). Furthermore, the ‘Contact Us’ module enables users to contact us and submit information on data not yet in cfMethDB. We are committed to processing new submissions and regularly update our database to reflect the latest findings.

## Discussion

In this study, we constructed cfMethDB, a comprehensive cfDNA methylation data resource for cancer biomarkers. We collected and analyzed cfDNA methylation data across various cancer types to identify potential biomarkers across the whole genome. cfMethDB provides not only search and analysis tools for users to discover biomarkers, but also comprehensive cfDNA fragment information, including fragment length and the 5’ end motif across different cancers.

In future versions, cfMethDB will be continually updated to include the following improvements:

(i) We will expand our data repository to encompass a wider range of cancer types. The aim of this expansion is to enhance the comprehensiveness and applicability of our database, thereby increasing its value for research and clinical applications.
(ii) We will collect single-molecule sequencing data to enrich our data resource. Single-molecule sequencing can simultaneously capture DNA sequence and DNA modification information without PCR amplification [48, 49]; single-molecule sequencing can also produce longer read than second-generation sequencing can produce [50]. Yu et al. demonstrated that methylation information derived from long reads sequencing can be used to determine the tissue origin of the fetal and maternal cfDNA with an area under the curve of 0.88 [51]. Although single-molecule sequencing data from cancer patients are currently limited, the potential of this approach should not be overlooked. In the future, we aim to incorporate single-molecule sequencing data to enhance the depth and breadth of our analyses.
(iii) We will introduce multimodal analysis module. cfDNA methylation data provides not only the methylation profile but also valuable fragment information, such as the 5’ end motif and fragment size. Based on our analysis of end motifs (Figure 4E, Figure S4), we infer that incorporating methylation status into the end motif information could enhance cancer diagnosis performance. However, a key challenge lies in the inherent differences between WGS and WGBS end motif analyses due to the bisulfite treatment used. Therefore, while WGBS end motif analysis provides valuable insights for further investigations, the results from WGS and WGBS should not be directly compared. In addition, we can identify genomic variants present in the cfDNA. Previous studies have shown that multimodal analysis of cfDNA can improve the performance of cancer diagnosis [52, 53]. In the future, we plan to integrate a multimodal diagnostic module into cfMethDB, with the goal of uncovering a broader spectrum of potential biomarkers through the synergistic analysis of diverse data types.
(iv) We will provide additional online functions based on user feedback. We are committed to keeping cfMethDB up to date to maintain its value as a user-friendly cfDNA methylation database. We hope that cfMethDB will facilitate the identification of cfDNA methylation biomarkers and contribute to the advancement of their clinical applications.

## Data availability

The cfMethDB database is available at https://cfmethdb.hzau.edu.cn/home and it can be accessed without registration or login.

## CRediT author statement

**Yuanhui Sun:** Data curation, Formal analysis, Visualization, Writing-review & editing. **Zhixian Zhu:** Data curation, Formal analysis, Writing-review & editing. **Qiangwei Zhou:** Formal analysis, Visualization, Writing-review & editing. **Zhe Wang:** Data curation. **Yuying Hou:** Data curation. **Xionghui Zhou:** Writing-review & editing. **Guoliang Li:** Conceptualization, Supervision, Writing-review & editing, Funding acquisition. All authors have read and approved the final manuscript.

## Supplementary material

Supplementary material is available at online

## Acknowledgments

We would like to thank Mr. Hao Liu from the National Key Laboratory of Crop Genetic Improvement for his essential assistance in managing the high-throughput computing clusters. We also appreciate the valuable feedback from the members of our research group on the database. This work was supported by the National Natural Science Foundation of China (Grant Nos. 32370630, 32400465, and 32250710678) and the National Key Research and Development Program of China (Grant No. 2021YFC2701201). The funders had no role in the study design, data collection and analysis, decision to publish, or preparation of the manuscript.

## References

[1] Bray F, Laversanne M, Sung H, Ferlay J, Siegel RL, Soerjomataram I, Jemal A. Global cancer statistics 2022: GLOBOCAN estimates of incidence and mortality worldwide for 36 cancers in 185 countries. CA Cancer J Clin 2024;74:229–63.

[2] Jamshidi A, Liu MC, Klein EA, Venn O, Hubbell E, Beausang JF, et al. Evaluation of cell-free DNA approaches for multi-cancer early detection. Cancer Cell 2022;40:1537-49.e12.

[3] Ginsburg O, Yip CH, Brooks A, Cabanes A, Caleffi M, Dunstan Yataco JA, et al. Breast cancer early detection: A phased approach to implementation. Cancer 2020;126 Suppl 10:2379–93.

[4] Gao Q, Lin YP, Li BS, Wang GQ, Dong LQ, Shen BY, et al. Unintrusive multi-cancer detection by circulating cell-free DNA methylation sequencing (THUNDER): development and independent validation studies. Ann Oncol 2023;34:486–95.

[5] Crosby D, Bhatia S, Brindle KM, Coussens LM, Dive C, Emberton M, et al. Early detection of cancer. Science 2022;375:eaay9040.

[6] Wan JCM, Massie C, Garcia-Corbacho J, Mouliere F, Brenton JD, Caldas C, et al. Liquid biopsies come of age: towards implementation of circulating tumour DNA. Nat Rev Cancer 2017;17:223–38.

[7] Ignatiadis M, Sledge GW, Jeffrey SS. Liquid biopsy enters the clinic -implementation issues and future challenges. Nat Rev Clin Oncol 2021;18:297–312.

[8] Jiang P, Chan CW, Chan KC, Cheng SH, Wong J, Wong VW, et al. Lengthening and shortening of plasma DNA in hepatocellular carcinoma patients. Proc Natl Acad Sci U S A 2015;112:E1317–25.

[9] Warren JD, Xiong W, Bunker AM, Vaughn CP, Furtado LV, Roberts WL, et al. Septin 9 methylated DNA is a sensitive and specific blood test for colorectal cancer. BMC Med 2011;9:133.

[10] Singh A, Gupta S, Badarukhiya JA, Sachan M. Detection of aberrant methylation of HOXA9 and HIC1 through multiplex MethyLight assay in serum DNA for the early detection of epithelial ovarian cancer. Int J Cancer 2020;147:1740–52.

[11] Ye J, Wu M, He L, Chen P, Liu H, Yang H. Glutathione-S-Transferase p1 Gene Promoter Methylation in Cell-Free DNA as a Diagnostic and Prognostic Tool for Prostate Cancer: A Systematic Review and Meta-Analysis. Int J Endocrinol 2023;2023:7279243.

[12] Li P, Liu S, Du L, Mohseni G, Zhang Y, Wang C. Liquid biopsies based on DNA methylation as biomarkers for the detection and prognosis of lung cancer. Clin Epigenetics 2022;14:118.

[13] Xu RH, Wei W, Krawczyk M, Wang W, Luo H, Flagg K, et al. Circulating tumour DNA methylation markers for diagnosis and prognosis of hepatocellular carcinoma. Nat Mater 2017;16:1155–61.

[14] Manoochehri M, Borhani N, Gerhäuser C, Assenov Y, Schönung M, Hielscher T, et al. DNA methylation biomarkers for noninvasive detection of triple-negative breast cancer using liquid biopsy. Int J Cancer 2023;152:1025–35.

[15] Yu F, Li K, Li S, Liu J, Zhang Y, Zhou M, et al. CFEA: a cell-free epigenome atlas in human diseases. Nucleic Acids Res 2020;48:D40–D4.

[16] Li M, Zhou T, Han M, Wang H, Bao P, Tao Y, et al. cfOmics: a cell-free multi-Omics database for diseases. Nucleic Acids Res 2024;52:D607–D21.

[17] Jiang P, Sun K, Peng W, Cheng SH, Ni M, Yeung PC, et al. Plasma DNA End-Motif Profiling as a Fragmentomic Marker in Cancer, Pregnancy, and Transplantation. Cancer Discov 2020;10:664–73.

[18] Liu Y. At the dawn: cell-free DNA fragmentomics and gene regulation. Br J Cancer 2022;126:379–90.

[19] Katz K, Shutov O, Lapoint R, Kimelman M, Brister JR, O’Sullivan C. The Sequence Read Archive: a decade more of explosive growth. Nucleic Acids Res 2022;50:D387–D90.

[20] Wang Y, Song F, Zhu J, Zhang S, Yang Y, Chen T, et al. GSA: Genome Sequence Archive. Genomics Proteomics Bioinformatics 2017;15:14–8.

[21] Chen S. Ultrafast one-pass FASTQ data preprocessing, quality control, and deduplication using fastp. Imeta 2023;2:e107.

[22] Felix Krueger FJ, Phil Ewels, Ebrahim Afyounian, Michael Weinstein, Benjamin Schuster-Boeckler, Gert Hulselmans, & sclamons. FelixKrueger/TrimGalore: v0.6.10. Zenodo 2023.

[23] Zhou Q, Lim JQ, Sung WK, Li G. An integrated package for bisulfite DNA methylation data analysis with Indel-sensitive mapping. BMC Bioinformatics 2019;20:47.

[24] Li H, Handsaker B, Wysoker A, Fennell T, Ruan J, Homer N, et al. The Sequence Alignment/Map format and SAMtools. Bioinformatics 2009;25:2078–9.

[25] Zhou Q, Guan P, Zhu Z, Cheng S, Zhou C, Wang H, et al. ASMdb: a comprehensive database for allele-specific DNA methylation in diverse organisms. Nucleic Acids Res 2022;50:D60–D71.

[26] Liu H, Liu X, Zhang S, Lv J, Li S, Shang S, et al. Systematic identification and annotation of human methylation marks based on bisulfite sequencing methylomes reveals distinct roles of cell type-specific hypomethylation in the regulation of cell identity genes. Nucleic Acids Res 2016;44:75–94.

[27] Yu G, Wang LG, He QY. ChIPseeker: an R/Bioconductor package for ChIP peak annotation, comparison and visualization. Bioinformatics 2015;31:2382–3.

[28] McKenna A, Hanna M, Banks E, Sivachenko A, Cibulskis K, Kernytsky A, et al. The Genome Analysis Toolkit: a MapReduce framework for analyzing next-generation DNA sequencing data. Genome Res 2010;20:1297–303.

[29] Quinlan AR, Hall IM. BEDTools: a flexible suite of utilities for comparing genomic features. Bioinformatics 2010;26:841–2.

[30] Moritz S, Bartz-Beielstein T. imputeTS: time series missing value imputation in R. R Journal 2017;9:207–18.

[31] Kuhn M. Building predictive models in R using the caret package. Journal of statistical software 2008;28:1–26.

[32] Buels R, Yao E, Diesh CM, Hayes RD, Munoz-Torres M, Helt G, et al. JBrowse: a dynamic web platform for genome visualization and analysis. Genome Biol 2016;17:66.

[33] Zhu Z, Zhou Q, Sun Y, Lai F, Wang Z, Hao Z, Li G. MethMarkerDB: a comprehensive cancer DNA methylation biomarker database. Nucleic Acids Res 2024;52:D1380–D92.

[34] Tang Z, Kang B, Li C, Chen T, Zhang Z. GEPIA2: an enhanced web server for large-scale expression profiling and interactive analysis. Nucleic Acids Res 2019;47:W556–W60.

[35] Mytilinaiou M, Nikitovic D, Berdiaki A, Kostouras A, Papoutsidakis A, Tsatsakis AM, Tzanakakis GN. Emerging roles of syndecan 2 in epithelial and mesenchymal cancer progression. IUBMB Life 2017;69:824–33.

[36] Essner JJ, Chen E, Ekker SC. Syndecan-2. Int J Biochem Cell Biol 2006;38:152–6.

[37] Oh T, Kim N, Moon Y, Kim MS, Hoehn BD, Park CH, et al. Genome-wide identification and validation of a novel methylation biomarker, SDC2, for blood-based detection of colorectal cancer. J Mol Diagn 2013;15:498–507.

[38] Wang L, Liu Y, Zhang D, Xiong X, Hao T, Zhong L, Zhao Y. Diagnostic accuracy of DNA-based SDC2 methylation test in colorectal cancer screening: a meta-analysis. BMC Gastroenterol 2022;22:314.

[39] Hu X, Wang Y, Zhang X, Li C, Zhang X, Yang D, et al. DNA methylation of HOX genes and its clinical implications in cancer. Exp Mol Pathol 2023;134:104871.

[40] Vega-Benedetti AF, Loi E, Moi L, Blois S, Fadda A, Antonelli M, et al. Clustered protocadherins methylation alterations in cancer. Clin Epigenetics 2019;11:100.

[41] Dong S, Li W, Wang L, Hu J, Song Y, Zhang B, et al. Histone-Related Genes Are Hypermethylated in Lung Cancer and Hypermethylated HIST1H4F Could Serve as a Pan-Cancer Biomarker. Cancer Res 2019;79:6101–12.

[42] Dong S, Lu Q, Xu P, Chen L, Duan X, Mao Z, et al. Hypermethylated PCDHGB7 as a universal cancer only marker and its application in early cervical cancer screening. Clin Transl Med 2021;11:e457.

[43] Dong S, Yang Z, Xu P, Zheng W, Zhang B, Fu F, et al. Mutually exclusive epigenetic modification on SIX6 with hypermethylation for precancerous stage and metastasis emergence tracing. Signal Transduct Target Ther 2022;7:208.

[44] Serpas L, Chan RWY, Jiang P, Ni M, Sun K, Rashidfarrokhi A, et al. Dnase1l3 deletion causes aberrations in length and end-motif frequencies in plasma DNA. Proc Natl Acad Sci U S A 2019;116:641–9.

[45] Chen M, Chan RWY, Cheung PPH, Ni M, Wong DKL, Zhou Z, et al. Fragmentomics of urinary cell-free DNA in nuclease knockout mouse models. PLoS Genet 2022;18:e1010262.

[46] Snyder MW, Kircher M, Hill AJ, Daza RM, Shendure J. Cell-free DNA Comprises an In Vivo Nucleosome Footprint that Informs Its Tissues-Of-Origin. Cell 2016;164:57–68.

[47] Hou Y, Meng XY, Zhou X. Systematically Evaluating Cell-Free DNA Fragmentation Patterns for Cancer Diagnosis and Enhanced Cancer Detection via Integrating Multiple Fragmentation Patterns. Adv Sci (Weinh) 2024;11:e2308243.

[48] Simpson JT, Workman RE, Zuzarte PC, David M, Dursi LJ, Timp W. Detecting DNA cytosine methylation using nanopore sequencing. Nat Methods 2017;14:407–10.

[49] Liu Q, Fang L, Yu G, Wang D, Xiao CL, Wang K. Detection of DNA base modifications by deep recurrent neural network on Oxford Nanopore sequencing data. Nat Commun 2019;10:2449.

[50] Amarasinghe SL, Su S, Dong X, Zappia L, Ritchie ME, Gouil Q. Opportunities and challenges in long-read sequencing data analysis. Genome Biol 2020;21:30.

[51] Yu SCY, Jiang P, Peng W, Cheng SH, Cheung YTT, Tse OYO, et al. Single-molecule sequencing reveals a large population of long cell-free DNA molecules in maternal plasma. Proc Natl Acad Sci U S A 2021;118.

[52] Liu J, Dai L, Wang Q, Li C, Liu Z, Gong T, et al. Multimodal analysis of cfDNA methylomes for early detecting esophageal squamous cell carcinoma and precancerous lesions. Nat Commun 2024;15:3700.

[53] Bie F, Wang Z, Li Y, Guo W, Hong Y, Han T, et al. Multimodal analysis of cell-free DNA whole-methylome sequencing for cancer detection and localization. Nat Commun 2023;14:6042.

