## Supplementary Captions for "cfMethDB: a comprehensive cfDNA methylation data resource for cancer biomarkers"

### **Supplementary material**

**Figure S1 Landscape of the cfMethDB datasets.**

**A.** Number of covered CpGs for each sample with the coverage ≥2. **B.** Average cfDNA fragment size of cfDNA samples across different projects and cancer types.

**Figure S2 A high density of hypermethylated DMCs across various cancer types.**

**A.** Density distribution of hypermethylated DMCs on chromosome 2. The highlighted region corresponds to chr2:176 Mb–176.5 Mb within *HOXD* genes. **B.** Density distribution of hypermethylated DMCs on chromosome 3. The highlighted region corresponds to chr3:147 Mb–147.5 Mb within *ZIC1/ZIC4* genes. **C.** Density distribution of hypermethylated DMCs on chromosome 5. The highlighted region corresponds to chr5:141 Mb–141.5 Mb within *PCDH* genes. **D.** Density distribution of hypermethylated DMCs on chromosome 7. The highlighted region corresponds to chr7:27 Mb–27.5 Mb within *HOXA* genes.

**Figure S3 Pan-cancer DNA methylation biomarker genes.**

**A.** Distribution of DMCs in the *SIX6* gene in various cancers. **B.** Distribution of DMCs in the *SOX11* gene in various cancers.

**Figure S4 Landscape of end motifs in cfMethDB datasets.**

**A.** Motif diversity score of cancer and normal samples across the whole genome with methylation status. End Motif *: 625 end motifs with methylation status ('U' represents unmethylated cytosines). **B.** Motif diversity score of cancer and normal samples across the whole genome without methylation status. End Motif: 256 end motifs without methylation status. P values are calculated by Wilcoxon rank sum test.

**Figure S5 Detailed sample information.**

**A.** Detailed information for sample GSM4502212. **B.** Number of cytosines at different methylation levels. **C.** Number of cytosines at different coverage levels and their percentage relative to the total number of covered cytosines. **D.** Number of reads at different fragment sizes and their percentage relative to the total number of reads. **E.** Coverage (upper panel) and effective sequencing depth (lower panel) for each chromosome (very few targeted regions are located on chromosome X and chromosome Y).

**Table S1 Summary of early detection kits for cancer approved by the NMPA.**

**Table S2 Statistics of cfDNA methylation data in cfMethDB.**

**Table S3 Statistics of DMCs in each cancer type.**

**Table S4 Summary of** **cfDNA methylation data in functional modules.**
