## Supplementary figures and images for "cfMethDB: a comprehensive cfDNA methylation data resource for cancer biomarkers"

### Figure S1

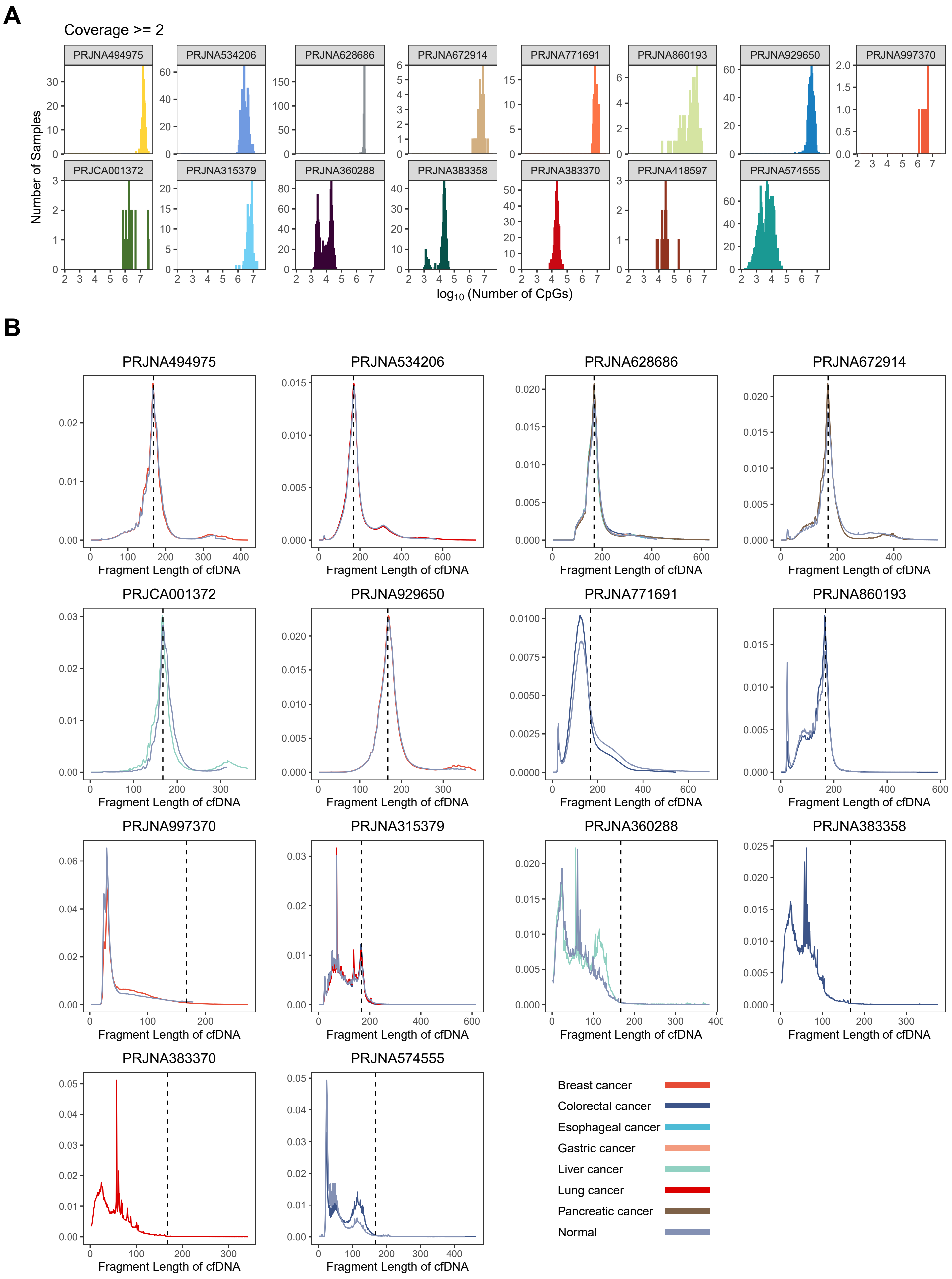

### Figure S2

***Figure S2***

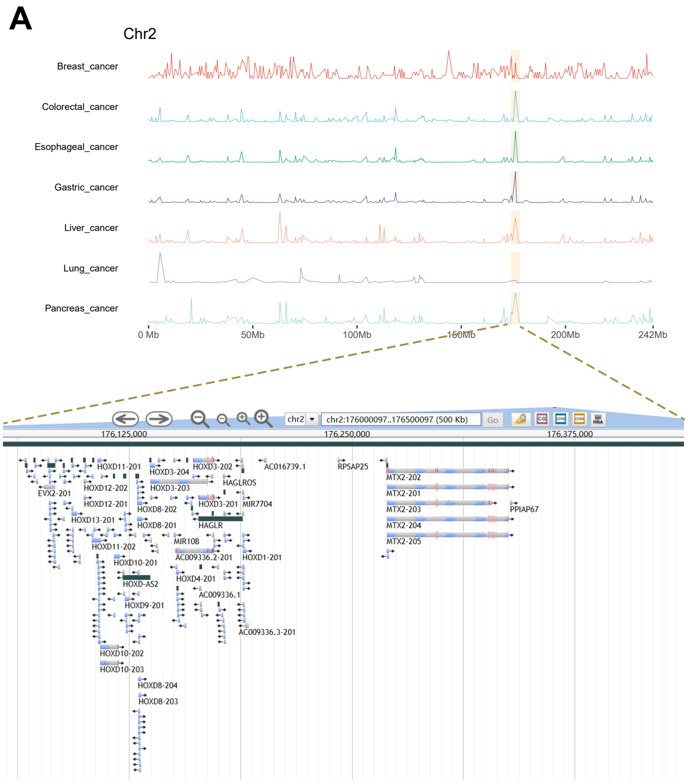


*HOXD Genes (chr2 : 176 Mb - 176.5 Mb)*

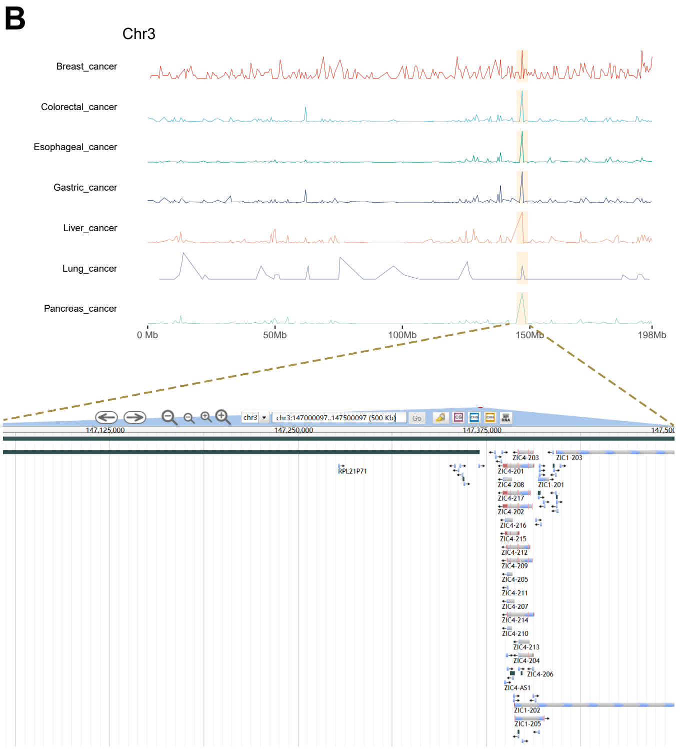


*ZIC1/ZIC4 Genes (chr3 : 147 Mb - 147.5 Mb)*

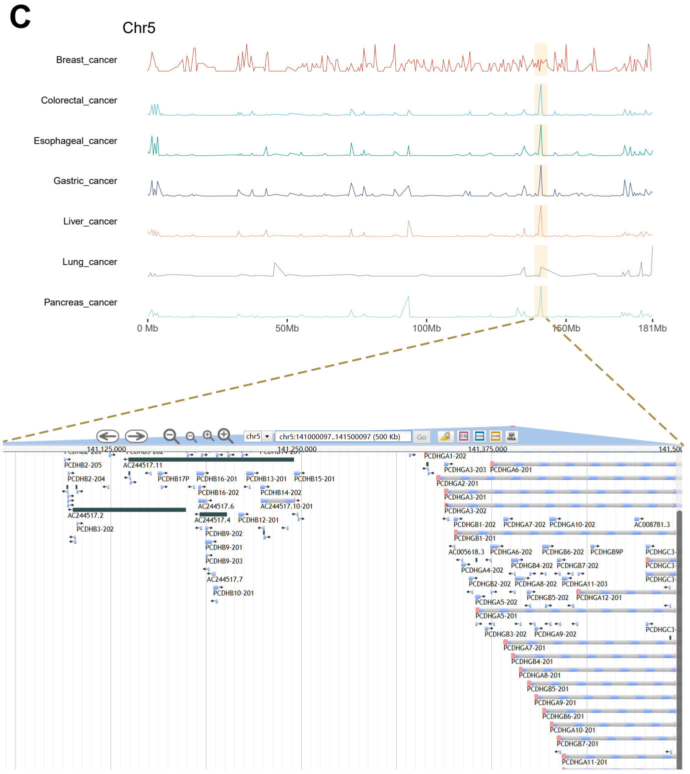


*PCDH Genes (chr5 : 141 Mb - 141.5 Mb)*

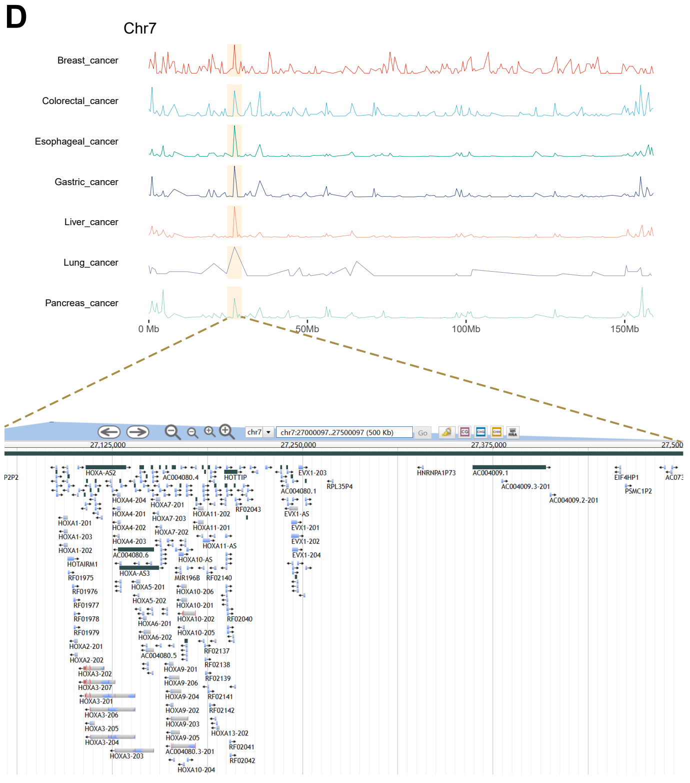


*HOXA Genes (chr7 : 27 Mb - 27.5 Mb)*

### Figure S3

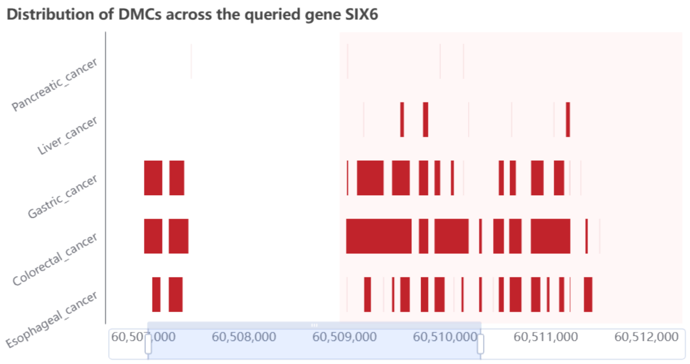

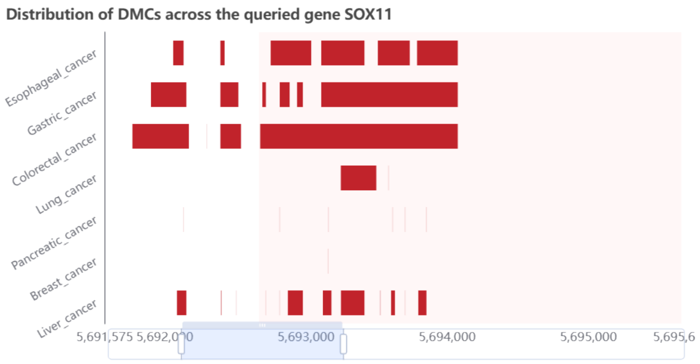

### Figure S4

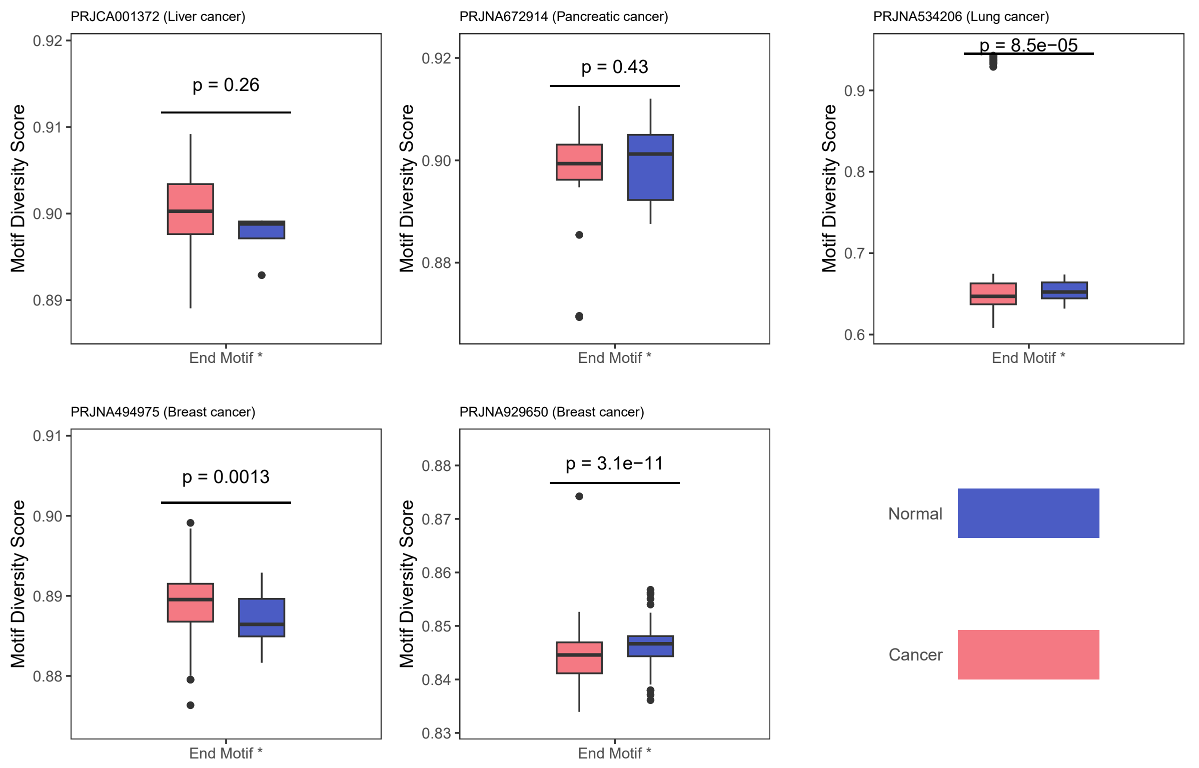

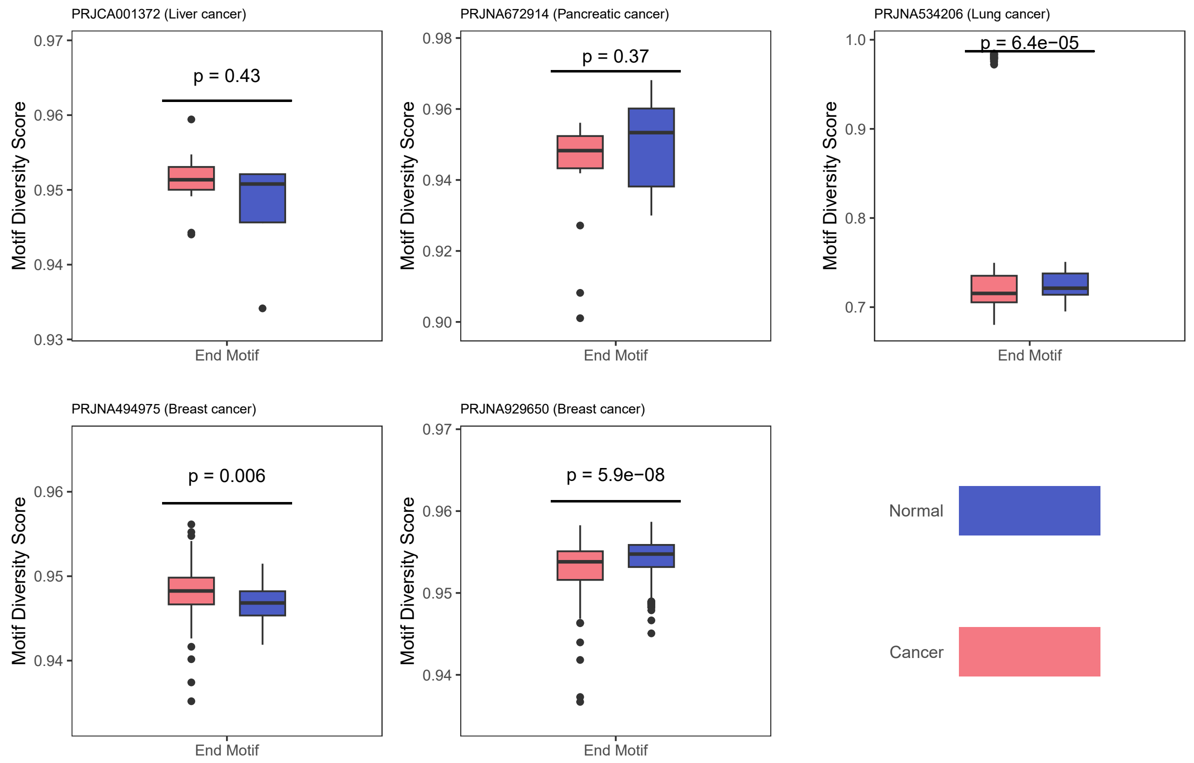
