## Supplementary material for "cfMethDB: a comprehensive cfDNA methylation data resource for cancer biomarkers": Figure S5

**A**

| BW | GEO ID | Sample | Study ID | Bases | Instrument | Layout | RUN | Title | Source | Public | Pubmed | bsrate |
|---|---|---|---|---|---|---|---|---|---|---|---|---|
| | GSM4502212 | Esophageal cancer | PRJNA628686 | 3.88Gb | Illumina Nova Seq 6000 | PAIRED | SRR11615982 | KRp2-102 | Plasma | 2020-04-30 | 34465601 | 0.9831 |

| Item | Value |
|---|---|
| GEO | GSM4502212 |
| Sample | Esophageal_cancer |
| Title | KRp2-102 |
| Organism | Homo sapiens |
| Instrument | Illumina NovaSeq 6000 |
| Source | Plasma |
| SRA | SRR11615982 |
| Bases | 3.88Gb |
| Public | 2020-04-30 00:00:00 |
| Layout | PAIRED |
| Library | Bisulfite-Seq |
| Pubmed | 34465601 |
| Tissue | Plasma |
| FTP | ftp.sra.ebi.ac.uk/vol1/fastq/SRR116/082/SRR11615982/SRR11615982_1.fastq.gz<br>ftp.sra.ebi.ac.uk/vol1/fastq/SRR116/082/SRR11615982/SRR11615982_2.fastq.gz |

**B**

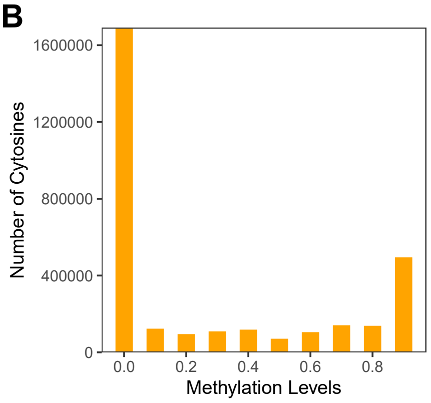


**C**

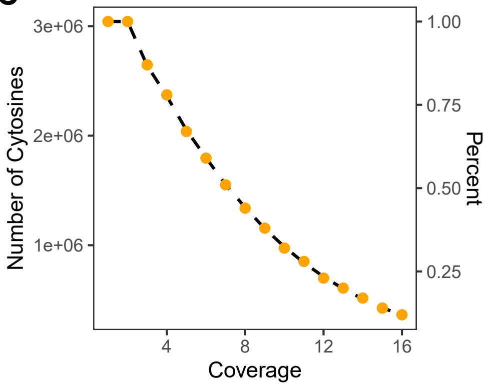


**D**

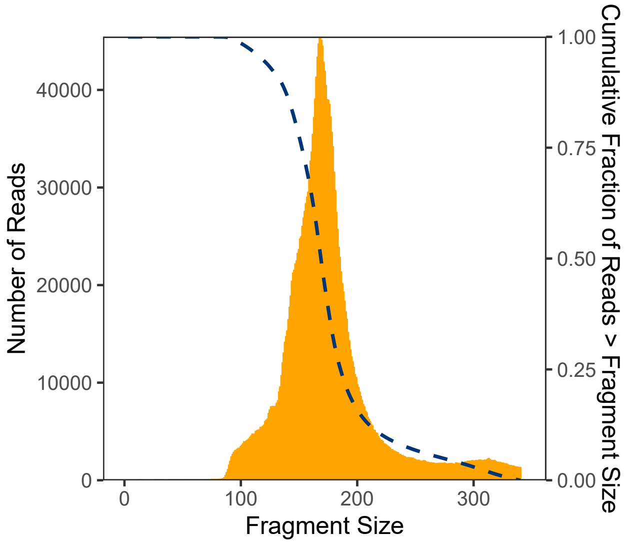


**E**

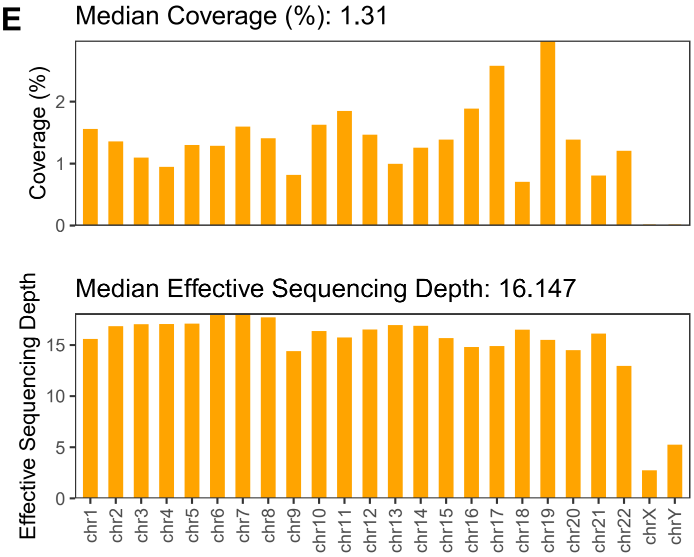
